# Time-resolved succession of epigenetic regulation during early mammalian development

**DOI:** 10.1101/503805

## Abstract

During early mammalian embryonic development, different epigenetic mechanisms undergo dramatic changes; yet how these interconnected epigenetic layers function together to coordinate expression of the genetic code in a spatiotemporal manner remains unknown. Here, we describe a time-resolved study of the hierarchy of epigenetic marks and events, which we used to model transcriptional programs that cannot be understood by investigating steady state. We found that, following fertilization, the re-establishment of accessible chromatin together with transcription factors inherited from oocytes initiates transcription at the 2-cell stage, and then the introduction of active histone modification, H3K4me3, facilitates gene up-regulation at the 4-cell stage, then stabilization of higher-order chromatin structures at the 8-cell stage further enhances transcriptional activity. During the first lineage specification at ICM, transcription activity multifaceted regulation of epigenetic marks. Finally, we quantitatively model the stage succession of different epigenetic marks on transcriptional programs during early embryonic development.

## Introduction

During early mammalian embryonic development, the transcriptional program is controlled by dynamic regulation of the epigenome. Interplay among different layers of epigenetic marks and their combinatorial effects contribute to major events in early embryonic development, such as zygotic gene activation (ZGA) [1, 2] and lineage commitment. The recent application of several advanced techniques has provided evidence documenting the mechanisms that mediate the transcriptional program. For example, the transition of the accessible chromatin landscape during mouse and human ZGA are both transcription-dependent [3]. Early cell-fate commitment is accompanied by extensive epigenetic reprogramming [4] and gene *Chaf1a* is essential for establishment of H3 lysine 9 trimethylation (H3K9me3) on long terminal repeats (LTRs) and subsequent transcriptional repression [5]. DNA methylation is crucial in many biological processes, including repression of gene transcription, maintenance of gene imprinting and X-chromosome inactivation, and repression of transposable elements [6–8].

Meanwhile, genome-wide studies of the epigenetic landscape have been used to observe the interplay between different marks. For example, the rebuilding of H3K9me3 on LTRs is involved in silencing their active transcription triggered by DNA demethylation [5]. A non-canonical form of histone H3 lysine 4 trimethylation (H3K4me3) in oocytes overlaps almost exclusively with partially methylated DNA domains [9]. Higher-order chromatin organization, which is associated with histone modifications [10, 11], undergoes drastic reprogramming after fertilization, followed by progressive maturation during early development [12]. Although recent technological advances provide us a comprehensive perspective of the dynamics of different epigenetic information during early mammalian development, integrative analyses are required to uncover the interconnections among different layers of epigenetic marks and further to document the spatiotemporal relationships between epigenetic regulation and transcriptional activity.

To address this gap, we performed a time-resolved analysis of transcriptional control during early embryonic development in mammals by integrating high-throughput sequencing data on chromatin accessibility, transcription factors (TFs), histone modifications, DNA methylation, and three-dimensional (3D) chromatin architecture. We investigated the relationship between gene expression and different epigenetic marks, and predicted the “writers” or “erasers” of transcriptional activity in 2-, 4-, and 8-cell embryos and the inner cell mass (ICM). Based on our findings, we provide a unifying picture for how epigenetic mechanisms regulate transcriptional activity in a stepwise manner.

## Results

### The 2-cell stage: Accessible chromatin initiates transcriptional activity

Following fertilization, several transcription factors (TFs), known as “pioneer” TFs recognize and bind to repressed chromatin and initiate events that lead to chromatin opening and binding of other TFs and epigenetic modifiers, thereby driving zygotic gene expression [1, 13–15]. To further understand the effects of the chromatin state and TFs on the regulation of gene expression, we measured the dynamics of chromatin accessibility and TF binding sites (TFBSs). Because RNA transcripts can be inherited from oocytes, we only analyzed genes that were activated in preimplantation embryos (denoted hereafter as “ZGA-only genes”, Supplementary Table 1).

Correlation analysis indicated that gene expression was significantly correlated with the density of accessible chromatin in promoters since 2-cell stage, when the major ZGA event occurs. The greatest correlation was observed in the 8-cell stage; thereafter, the correlation underwent a sharp decrease in the ICM and mouse embryonic stem cell (mESC) stages (Fig. 1A). The protamine-packaged sperm genome undergoes dramatic remodeling, and chromatin accessibility of the paternal genome becomes largely reprogrammed by the PN3 stage [16, 17]. We found that global accessible chromatin was widely established by the 2-cell stage, and over 20% of the accessible chromatin was in the promoter region (Fig. 1B). Moreover, much of accessible chromatin that were maintained from previous stage were close to the gene transcription start sites (TSSs) (Supplementary Fig. 1A), allowing the transcriptional machinery to access gene promoters. Many key genes were accessible at the 2-cell stage or earlier. For example, *Arntl*, which is associated with the rhythmic opening of chromatin at promoters and regulates DNA accessibility for other TFs [18], was accessible during the entire preimplantation stage. *Eif2b2*, a translation initiation factor, and *Nfya*, which forms a histone-like structure that binds to DNA and promotes chromatin accessibility and ZGA [13, 16, 19], were accessible from the 2-cell embryo stage (Fig. 1C).

**Fig. 1.**
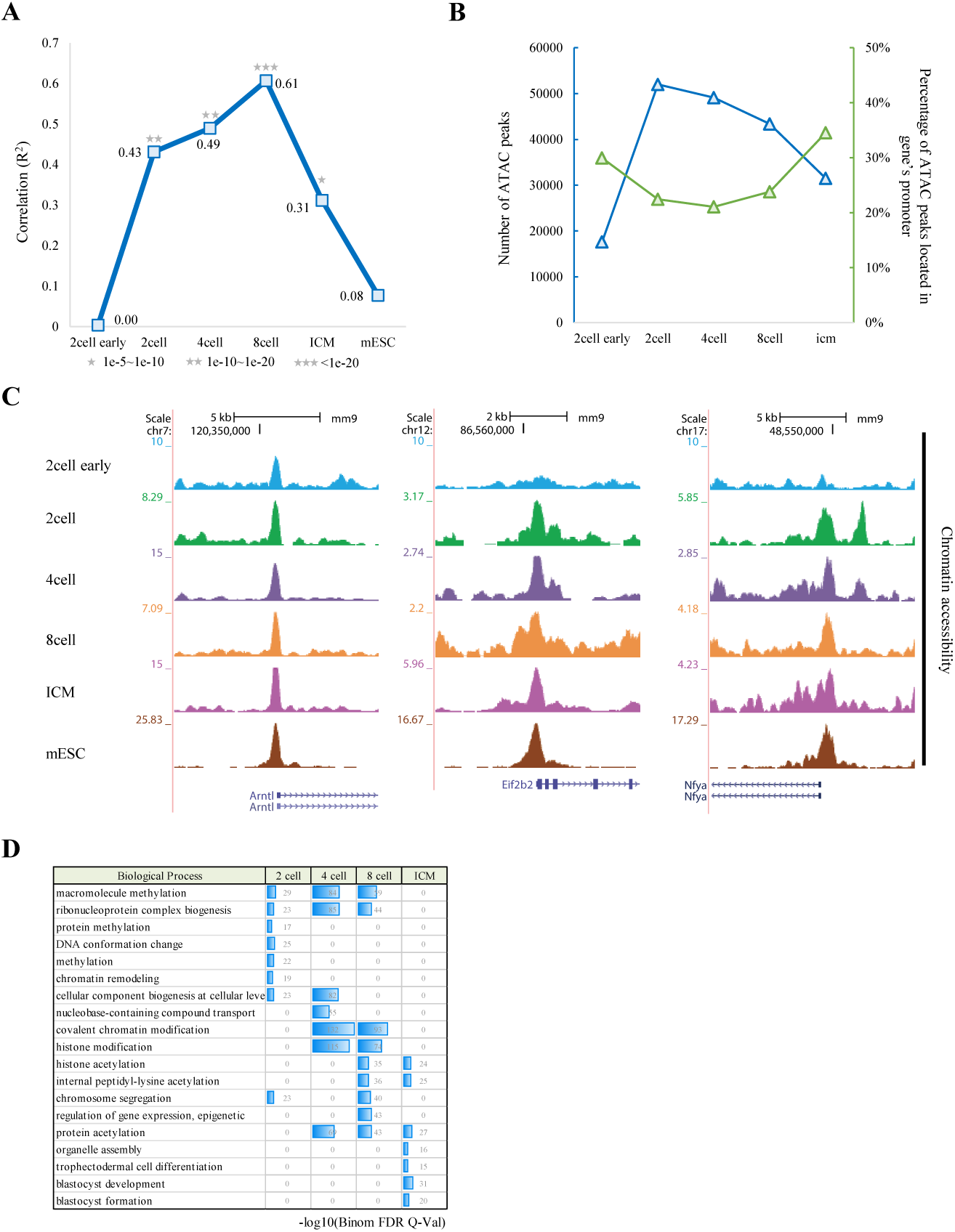
General characterization of accessible chromatin during early embryonic development. **A**. Correlations between gene expression and chromatin accessibility in promoters (2 kb upstream of TSSs) of ZGA-only genes. Correlation coefficients (R^2^ values) were calculated by Pearson’s linear correlation. Significance is indicated by grey stars. **B**. Basic information on accessible chromatin during early embryonic development. Blue line shows number of ATAC peaks. Green line shows percentage of ATAC peaks in gene promoter (2 kb upstream of TSS). **C**. UCSC plots showing chromatin accessibility in promoters of genes *Arntl, Eif2b2*, and *Nfya*. **D**. Gene Ontology terms for genes associated with ATAC peaks from previous stages and corresponding *p*-values.

Once established, accessible chromatin tends to be maintained [16]. Thus, we investigated the potential function of the maintained accessible chromatin by applying the program GREAT [20] (Fig. 1D). Accessible chromatin maintained from the early to late 2-cell embryo stage was associated with chromatin conformation and methylation, whereas there was a strong relationship between chromatin accessibility and histone modification in 4- and 8-cell embryos, indicating a potential role of accessible chromatin in regulating histone modifications, and in 8-cell embryo, maintained accessible chromatin was also associated with chromosome segregation, indicating a regulatory role in chromatin structure by accessible chromatin. In the ICM, accessible chromatin was enriched for lineage specification, including trophectodermal cell differentiation and blastocyst development, suggesting that the chromatin regulatory landscape had been prepared for the first-cell lineage specification [16].

Because accessible chromatin is often occupied by TFs, we next identified TFBSs at gene promoter regions. In general, we found a very low correlation between TFBSs density and expression of ZGA-only genes in early 2-cell (Fig. 2A) and Only 23% (364/1582) of ZGA-only genes were accessible (Supplementary Table 2). One possible explanation for this result is that accessible chromatin levels are difficult to measure by the assay for transposase-accessible chromatin followed by sequencing (ATAC-seq) in early 2-cell embryos. Another possibility is that genes in the very early stage are self-initiating and may not depend on accessible chromatin. The relationship between TFBSs and gene expression was highest in the 4- and 8-cell stages but sharply decreased in the ICM and mESCs (Fig. 2A). We then examined the expression levels of TF genes. Interestingly, dynamic changes in the expression levels of TF genes were consistent with the correlation reported above (Fig. 2B). A possible explanation for this finding is that adequate amounts of TF RNA transcripts were required for the binding of TFs to DNA sequences. To test this hypothesis, we further investigated the TF binding data by chromatin immunoprecipitation sequencing (ChIP-seq) in a well-characterized human embryonic stem (ES) cell line. TFs with sufficient binding sites in the accessible chromatin and adequate RNA transcripts, such as Sp1 and Sp4, were detected by ChIP-seq, whereas TFs with insufficient binding sites or inadequate RNA transcripts, such as Sp2 and Srf, were not clearly detected (Fig. 2C).

**Fig. 2.**
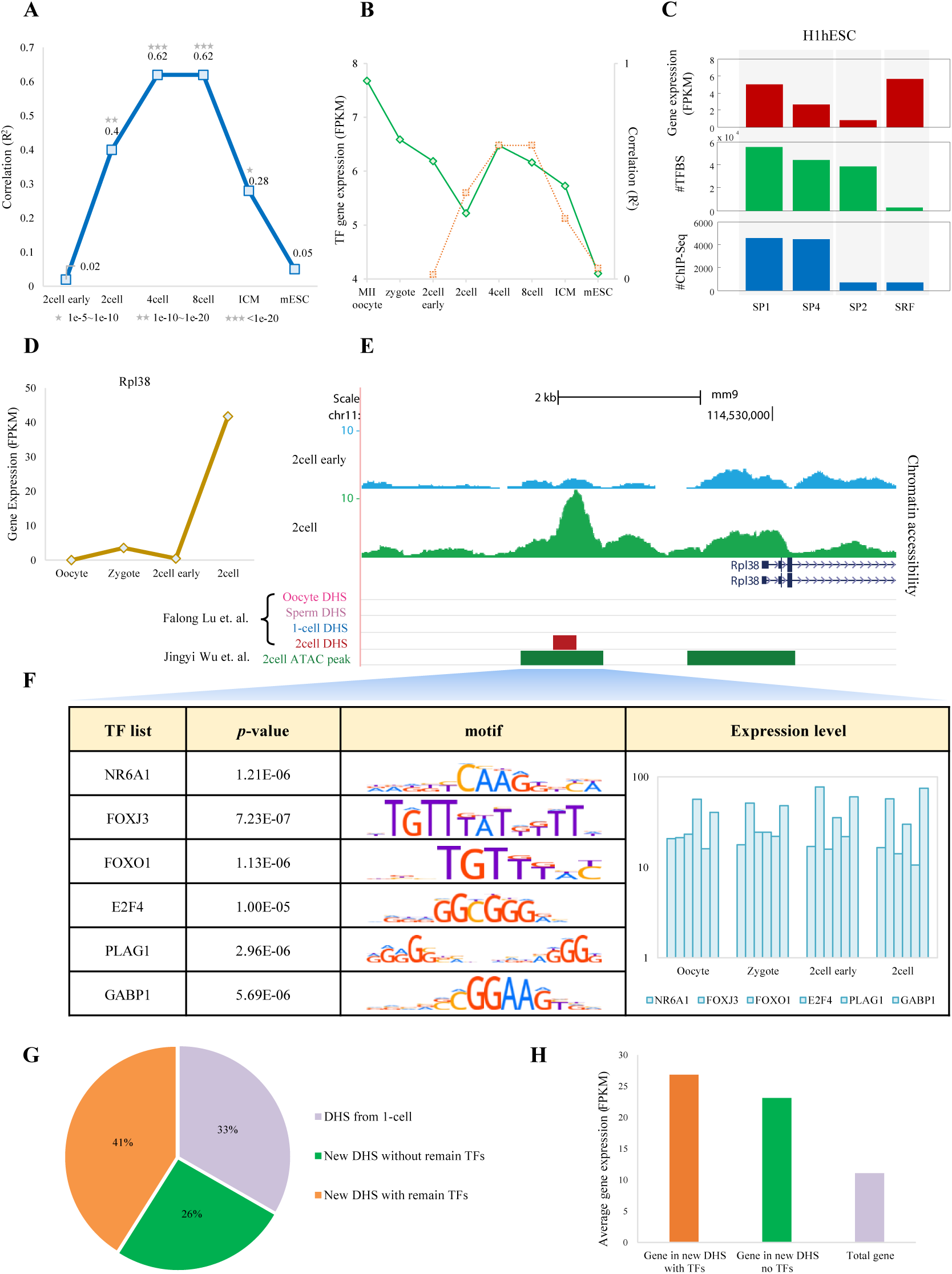
TFs bind to accessible chromatin and further initiate transcriptional programs. **A**. Correlations between gene expression and TFBS density in promoter regions (2 kb upstream of TSSs) of ZGA-only genes. Correlation coefficients (R^2^ values) were calculated by Pearson’s linear correlation. Significance is indicated by grey stars. **B**. Blue line showed average expression level for 395 TFs, determined as described in Methods, orange line showed correlation coefficients between gene expression and TFBS density. **C**. Red bar indicates expression levels of genes *SP1*, *SP2, SP4*, and *SRF*. Green bar indicates numbers of SP1-, SP2-, SP4-, and SRF-associated TFBSs scanned in promoters of all protein-coding genes. Blue bar shows numbers of SP1, SP2, SP4, and SRF ChIP-seq reads detected in promoters of all protein-coding genes. **D**. As an example of the activation of transcriptional programs by TFs in accessible chromatin, the gene *Rpl38* was silenced until the 2-cell stage. **E**. UCSC plot showing chromatin accessibility of *Rpl38* in the early 2-cell and 2-cell stages. DHSs and ATAC peaks by other studies are also shown. **F**. TFBSs scanned in corresponding ATAC peak. *p*-value is calculated by FIMO. Motif figures are collected from the HOCOMOCO (v10) database. Bar plot shows expression levels of TFs. **G**. Distribution of DHSs in the 2-cell embryo. Grey section includes DHSs detected in both 1- and 2-cell stages, whereas green and orange sections indicate DHSs only detected in the 2-cell stage. DHSs that can be targeted by highly expressed TFs (FPKM ≥ 10) are colored by orange and cannot be targeted by TFs with high expression level are colored by green. **H**. Expression level of genes associated with the three types of DHSs in G.

To understand how accessible chromatin affects activation of transcriptional programs, we characterized a class of active genes that were regulated by TFs inherited from oocytes (RPKM ≥ 10 from oocytes to 2-cell stage). For example, *Rpl38*, a gene related to RNA binding and a structural constituent of ribosome, was silenced until the 2-cell stage (Fig. 2D). We compared chromatin accessibility between the 2-cell and previous stages by integrating DNase I hypersensitivity (DHS) [16] and ATAC-seq data [21]. As expected, the promoter of *Rpl38* was accessible by the 2-cell stage (Fig. 2E). Several highly expressed TFs (RPKM ≥ 10), such as NR6A1, FOXJ3 and FOXO1, can bind to accessible chromatin, thus may contribute to the activation of *Rpl38* (Fig. 2F). Genome-wide analysis showed that two-thirds of the 2-cell DHSs were new from the 1-cell embryo stage. Of these DHSs, over 60% were accessible for highly expressed TFs (Fig. 2G). Genes with new DHSs targeted by TFs were expressed at higher levels than other genes (Fig. 2H). Together, these findings show that, in the 2-cell embryo, TFs inherited from oocytes bind to cz’s-regulatory elements embedded in accessible chromatin and may contribute to activation of transcriptional programs (Supplementary Fig. 1B).

### The 4-cell stage: H3K4me3 facilitates gene up-regulation

The dynamic process of chromatin accessibility is always accompanied by the redistribution of different histone marks [15, 22–24]. We next investigated the details and contribution of H3K4me3 dynamics to transcriptional activity. As H3K4me3 is detected in promoters and at low levels in distal regulatory elements [25], we examined the density of H3K4me3 in promoters of ZGA-only genes. Generally, gene expression was weakly correlated with H3K4me3 density in the early 2-cell and 2-cell embryos, whereas the correlation sharply increased from the 4-cell stage (Fig. 3A).

**Fig. 3.**
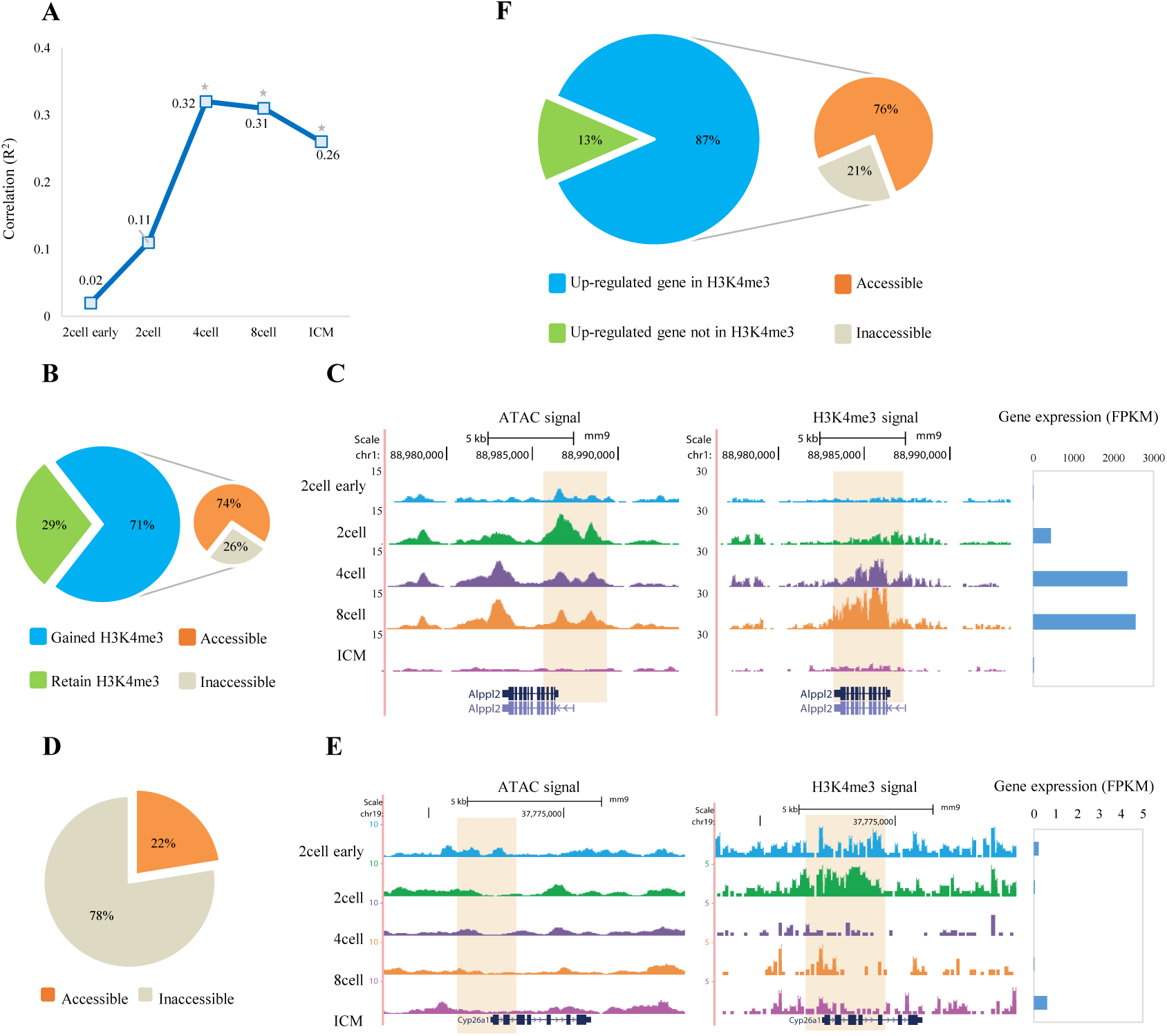
H3K4me3 facilitates gene up-regulation in the 4-cell stage. **A**. Correlations between gene expression and H3K4me3 signal levels in promoters (2 kb upstream of TSSs) of ZGA-only genes. Correlation coefficients (R^2^ values) were calculated by Pearson’s linear correlation. Significance is shown as grey stars. **B**. Pie charts showing (left) the percentage of gained H3K4me3 peaks in the 4-cell stage and (right) the percentage of gained H3K4me3 that are accessible. **C**. UCSC plots showing chromatin accessibility and H3K4me3 signal surrounding gene *Alppl2*. Bar plot showing expression level of gene *Alppl2*. **D**. Pie chart showing the percentage of lost H3K4me3 peaks that are accessible in 4-cell stage. **E**. UCSC plots showing chromatin accessibility and H3K4me3 signal surrounding gene *Cyp26a1*. Bar plot showing expression level of gene *Cyp26a1*. **F**. Pie charts showing (left) the percentage of up-regulated genes with and without H3K4me3 peaks in the promoter and (right) chromatin accessibility of up-regulated genes that are targeted by H3K4me3 peaks in the promoter.

To reveal the relationship between the dynamics of H3K4me3 and chromatin accessibility and their combinatory effects on transcriptional activity, we analyzed the gain or loss of H3K4me3 peaks in 4-cell embryos. Over 70% of H3K4me3 peaks in the 4-cell embryo were new compared to H3K4me3 peaks in the 2-cell embryo, and almost 75% of these gained H3K4me3 peaks was accessible in the 4-cell or earlier stages (Fig. 3B). For example, the promoter of *Alppl2* was accessible in the 2-cell embryo, and the H3K4me3 level increased in the 4- and 8-cell stages, as a result, *Alppl2* was active in the 2-cell stage, and its expression level increased nearly 6-fold in the 4- and 8-cell stages. However, *Alppl2* was silenced in the ICM, which we attributed to the closed chromatin and low H3K4me3 signal (Fig. 3C). Next, we focused on the H3K4me3 peaks that disappears from the 4-cell stage. Interestingly, over 75% of the lost H3K4me3 peaks was inaccessible before the 4-cell stage (Fig. 3D). For example, the promoter of *Cyp26a1* was inaccessible during early embryonic development. The H3K4me3 peak was present in the early 2-cell and 2-cell stages but disappeared thereafter. As a result, *Cyp26a1* was silenced during the entire implantation period (Fig. 3E).

To summarize, a strong correlation between H3K4me3 level and gene expression was observed from the 4-cell stage on. In the 4-cell embryos, 87% of all up-regulated genes were targeted by H3K4me3 in the promoters, and over 75% of these genes were accessible (Fig. 3F). Gain and loss of the H3K4me3 signal were directly associated with chromatin accessibility and they may together facilitated gene up-regulation (Supplementary Fig. 2A).

### The 8-cell stage: 3D chromatin structures enhance transcriptional activity

Recent studies reported that histone marks, eRNA, and chromatin accessibility are involved in chromatin organization [26–29]. Therefore, we examined the establishment of higher-order chromatin structure and lower-order chromatin loops and their effects on gene expression. Dynamic analysis showed 37% to 53% of the topologically associating domain (TAD) boundaries were new in each stage from the 2- to 8-cell stage, whereas fewer than 30% of TAD boundaries were new after the 8-cell stage (Fig. 4A). Gene TSSs were significantly located in TAD boundaries (*p*-value < 0.01, permutation test; Supplementary Fig. 3A). Expression levels of genes located in TAD boundaries were significantly higher than those outside TAD boundaries since 2-cell stage and the express level reached its highest in 8-cell stage (Fig. 4B). These results indicated that chromatin structures undergoes dramatic changes before the 8-cell stage and associated with transcription activity since then.

**Fig. 4.**
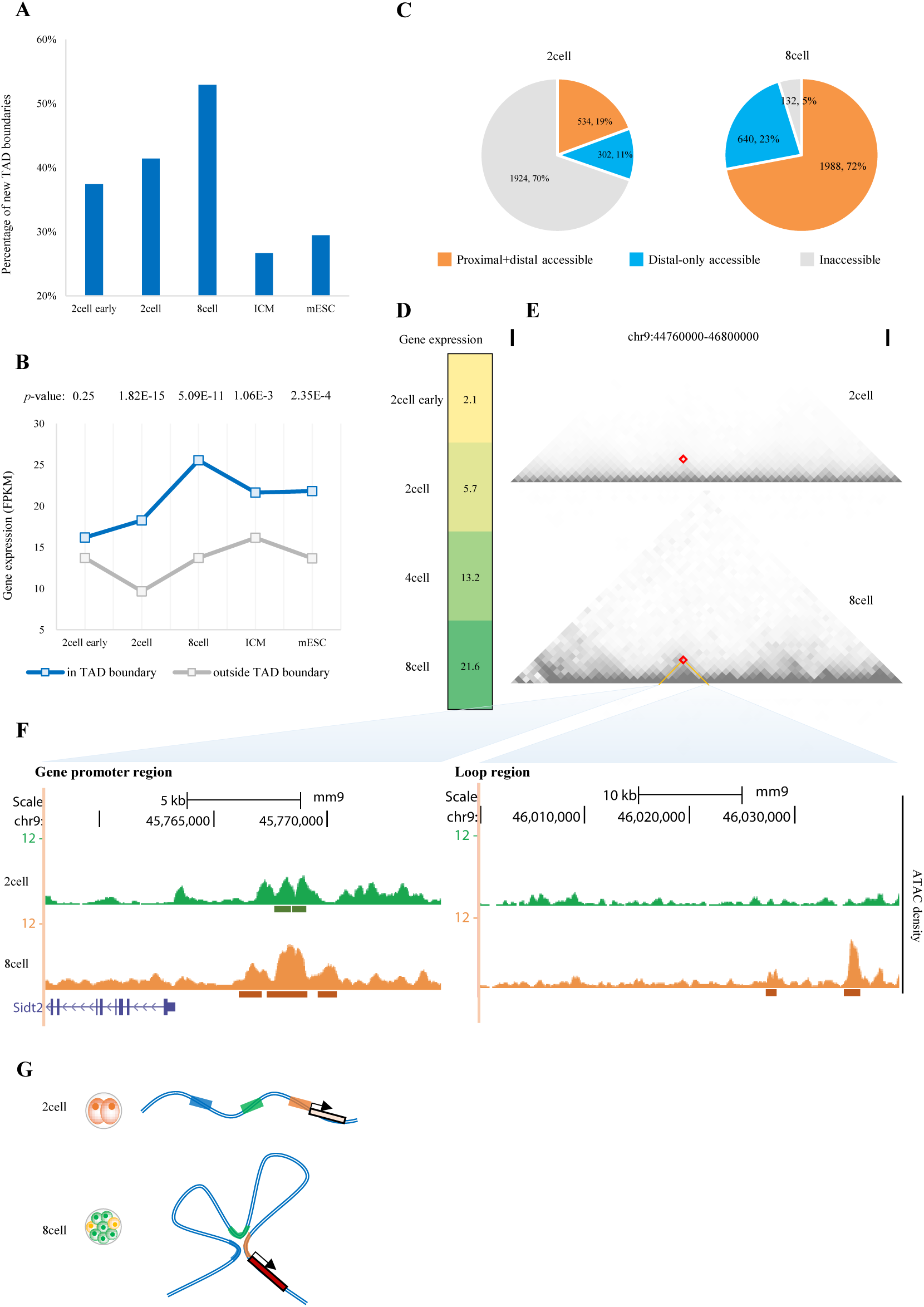
3D chromatin structures enhance transcriptional activity in the 8-cell stage. **A**. Bar plot showing the percentage of TAD boundaries that are new in the current compared to the previous stage. **B**. Average expression level of genes located in TADboundary and outside TAD boundary. P-value were calculated by T test. **C**. Pie charts showing genes with accessible chromatin in the promoter or in the chromatin interaction loop region. **D**. Expression level of gene *Sidt2*. **E**. Hi-C heatmaps of gene *Sidt2* and associated loop region at 40-kb resolution for 2- and 8-cell embryos. **F**. UCSC plots showing chromatin accessibility of gene *Sidt2* and associated loop region. **G**. Model showing that accessible chromatin in the 8-cell stage is condensed by widespread interaction loops, thus enhancing transcriptional activity.

We next explored the effect of chromatin loops on gene regulation. The number of Hi-C significant interactions increased after fertilization (Supplementary Fig. 3B), indicating a complex chromatin organization that was compressed into densely connected loops. As open chromatin regions direct the binding of key TFs (e.g., CTCF [29, 30]) and subsequently mediate chromatin looping; therefore, we compared the accessibility of chromatin loops in the 2- and 8-cell stages (4-cell Hi-C data were not available). Over 50% of loop-spanned regions containing accessible chromatin in the 8-cell embryo, whereas only 20% in the 2-cell embryo (Supplementary Fig. 3C). Meanwhile, 70% of up-regulated genes were inaccessible at distal regions in the 2-cell stage, indicating weak regulation by chromatin looping. However, 95% of up-regulated genes in the 8-cell stage were accessible at the promoter or distal chromatin interaction loops and, thus, can be targeted by regulatory factors such as TFs (Fig. 4C).

Chromatin accessibility of spatially proximate promoter and distal regions generally contributed to the increase of gene expression. Therefore, we examined the chromatin accessibility and Hi-C interactions at genes that were up-regulated in the 8-cell stage. For example, *Sidt2*, which mediates direct uptake of DNA [31], was active in the 2-cell stage and up-regulated in the 8-cell stage (Fig. 4D). No loop was detected surrounding *Sidt2* in the 2-cell stage, but a significant interaction (*p*-value = 2.9e-38) was observed in the 8-cell stage (Fig. 4E). The promoter of *Sidt2* was accessible in both stages, but only the interaction region in the 8-cell stage was open (Fig. 4F). By contrast, *Gnl2* was inaccessible in the promoter, but the expression was up-regulated in the 8-cell stage (Supplementary Fig. 3D, F). We observed a significant interaction (*p*-value = 1.5e-19) at the region 130 kb upstream of *Gnl2*, which was highly accessible (Supplementary Fig. 3E, F).

To summarize, these findings show that TADs tend to be consolidated as late as the 8-cell stage, consistent with a previous report [32], and that TAD boundaries constrain high gene expression. Compared to the 2-cell stage, accessible chromatin in the 8-cell stage is condensed by widespread interaction loops, thus enhancing transcriptional activity (Fig. 4G).

### The ICM: Multifaceted regulation of epigenetic marks

The inner cell mass (ICM) and the trophectoderm (TE) begin to segregate when the first lineage specification starts at the morula stage [33]. Dramatic changes of epigenetic marks occur at that time, H3K27me3 begins to emerge at canonical Polycomb target promoters [34] and the signal were significant higher than that at the onset of zygotic transcription in 2-cell embryos (Supplementary Fig. 4A). DNA demethylation is completed and then sharply increased post-implantation [35, 36] (Supplementary Fig. 4B), the CG methylation of gene’s promoter was much weak in ICM compared to that in 2-cell stage (Supplementary Fig. 4C). Large-scale H3K9me3 reestablishment occur immediately after fertilization and the disequilibrium of parental H3K9me3 lasted until the ICM stage [37]. Also, the imbalance in parental H3K4me3 signals lasts until ICM [38] but the global signal of H3K4me3 in ICM was comparable to that in 2-cell stage (Supplementary Fig. 4D). Thus, the transcription program multifaceted the regulation of epigenetic marks in ICM.

Correlation analysis showed that gene expression was significant positive correlated with chromatin accessibility and H3K4me3 level in gene’s promoter while significant negative correlated with H3K27me3 (Fig. 5A). H3K9me3 was weak correlated with gene expression. For DNA methylation, we observed a strong correlation between mCG/CG level and gene expression, but the mCG/CG level was sharply decreased to 0.05 when expression level (FPKM) > 3 (Fig. 5A). We further examined the epigenetic signals surrounding gene’s TSS (Fig. 5B), ATAC-seq signal and H3K4me3 signal were enriched in genes with high expression level while broad H3K27me3 domains were enriched in silence genes. However, H3K9me3 signal and mCG/CG level were indiscriminate. In addition, strong correlations were observed among chromatin accessibility, H3K4me3, H3K27me3 and CG methylation, but H3K9me3 was weak correlated with other markers (Supplementary Fig. 4E).

**Fig. 5.**
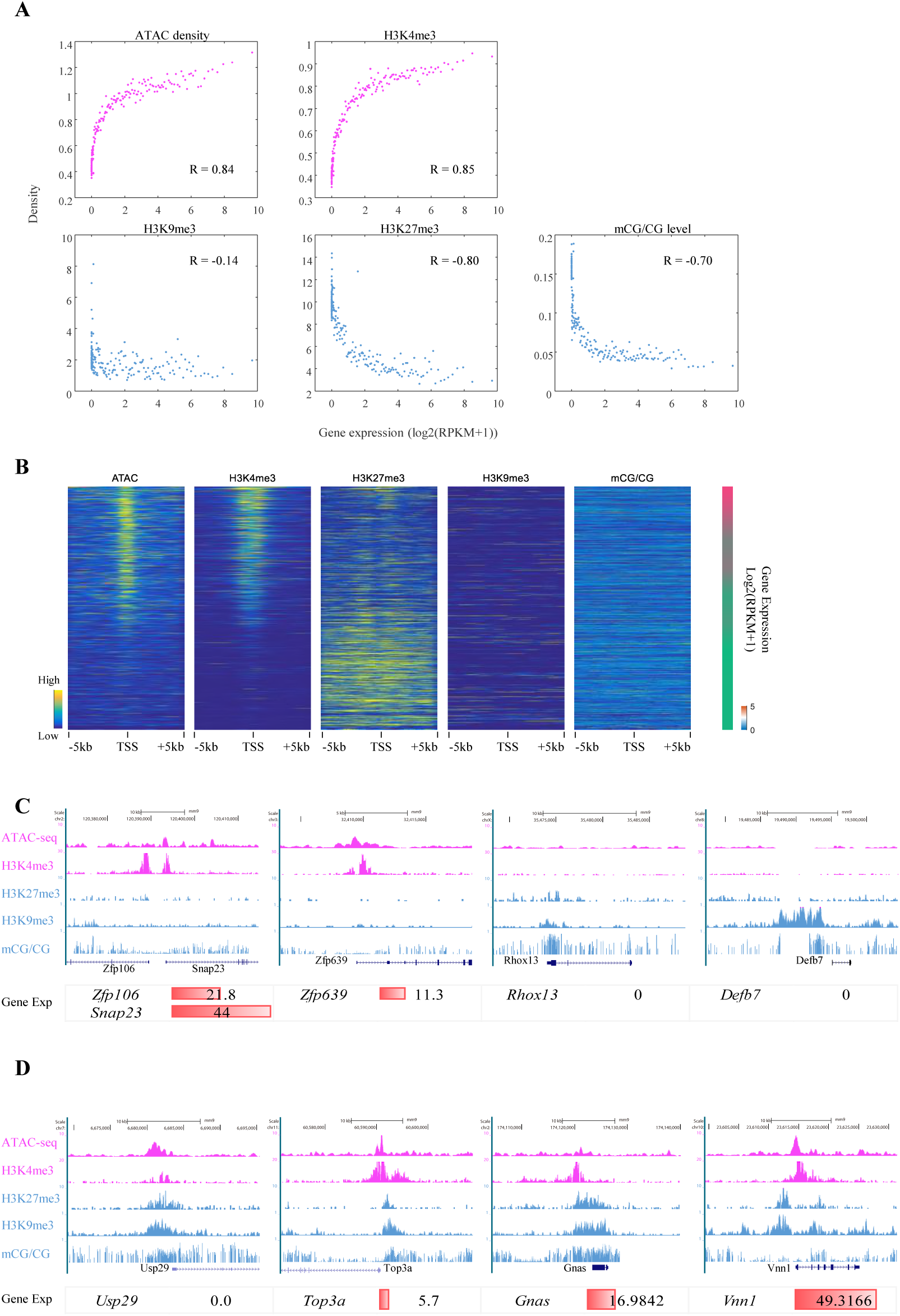
Multifaceted regulation of epigenetic marks at ICMs. **A**. Scatter plots showing the correlations between gene expression and epigenetic signals in promoters (2 kb upstream of TSSs). Correlation coefficients were calculated by Pearson’s linear correlation. **B**. Heatmaps showing the enrichment of epigenetic marks (normalized RPKM) in ICM stage, bar chart showing the expression level of corresponding genes. **C, D**. UCSC plots showing epigenetic signals surrounding gene *Zfp106, Snap23, Zfp639, Rhox13, Defb7, Usp29, Top3a, Gnas* and *Vnn1*. Bar plot shows gene expression level.

To illustrate the multifaceted regulation of epigenetic marks at ICM, we focus on genes with strong epigenetic signals. Genes targeted by strong active markers but weak poised markers were highly expressed while the expression level of genes targeted by strong poised markers but weak active markers were very low (Fig. 5C). In addition, when genes were targeted by both strong active and poised markers, the transcription activity was indeterminable (Fig. 5D). Furthermore, we investigated all active genes (FPKM > 10, 3862) and silence genes (FPKM < 1, 10555) in ICM (Supplementary Fig. 5A). 26% of active genes were targeted by both strong accessible chromatin and H3K4me3 signal in the promoter and 38% were targeted by either strong accessible chromatin or H3K4me3 signal. Meanwhile, about 43% of silence genes were targeted by at least one poised marker, but rare silence genes (less than 1%) were targeted by all these three poised markers in the promoter.

To summarize, in ICM, Major modifications will together contribute to the regulation of gene expression in ICM as accessible chromatin and H3K4me3 perform as an activator and H3K9me3, H3K27me3 and DNA methylation perform as a strong barrier of transcriptional program.

### Different layers of epigenetic marks shape transcriptional programs

Recent studies provide insights into the dynamics of epigenetics during early embryonic development, but much remains unknown about how different epigenetic marks control expression of the genetic code in a stepwise manner. To this end, we integrated different layers of epigenetic marks and quantitatively described their effects on transcriptional activity. Our analysis was conducted on ZGA-only genes, which are activated in preimplantation embryos (Supplementary Fig. 6A). By performing linear regression, we first evaluated the regulation intensity for each epigenetic marks, including chromatin accessibility, active histone H3K4me3, chromatin structure and CG methylation (Fig. 6A). For missing data (e.g., Hi-C data in the 4-cell and DNA methylation in the 8-cell stage), we obtained the corresponding correlation by interpolation. In the early 2-cell stage, we observed weak levels of epigenetic regulation (Fig. 6A), both the genome-wide transcriptional activity (Supplementary Fig. 6A) and chromatin accessibility was general weak (Fig. 1B). However, a large amount of residual TF mRNA remained from the oocytes (Fig. 2B and Supplementary Fig. 6B), which may lead to chromatin opening and recruitment of chromatin-remodeling complexes. This process would facilitate binding of “reader”, “eraser”, and “writer” complexes, enabling different epigenetic marks to direct or facilitate gene expression. In the 2-cell stage, with the completion of paternal chromatin accessibility reprogramming [16], genome-wide accessible chromatin plays a major role in activating transcriptional programs and may lead to ZGA. In the 4-cell stage, there was an increased effect of the active histone modification H3K4me3, and a portion of genes targeted by H3K4me3 were activated or up-regulated. Further to the 8-cell stage, as spatial segregation of the chromatin structure can be found and the allelic differences in chromatin compartments are clear [32], higher-order chromatin architecture separated genes with high expression levels and chromatin interactions provided spatial regulation of gene expression. During the first lineage specification, cells segregated into the ICM and trophectoderm [4], and the process of transcriptional regulation is very complex, while multiple epigenetic marks together function in up- or down-regulating gene expression. Furthermore, we performed a multiple regression analysis to combine major epigenetic factors in regulating gene expression. As expected, after ZGA at 2-cell stage, the density of epigenetic marks is significant correlated with transcription activity (Supplementary Fig. 6C).

**Fig. 6.**
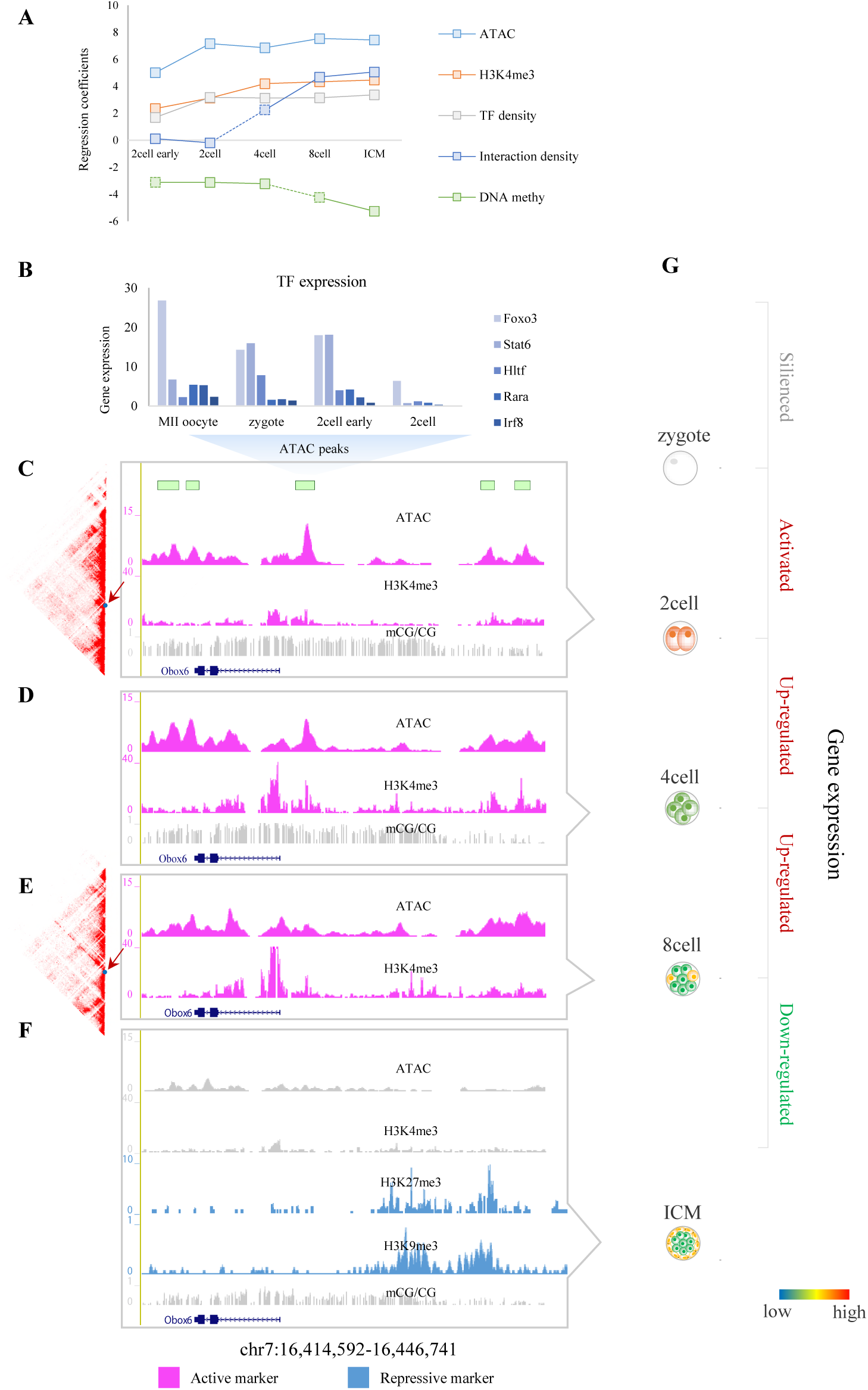
Different layers of epigenetic marks shape transcriptional programs. **A**. By performing linear regression, we quantitatively describe the effects of different layers of epigenetic marks on transcriptional activity in preimplantation embryos. This analysis was conducted on ZGA-only genes. **B-G**. Gene *Obox6* was used as an example to illustrate the relationship between different layers of epigenetic information and transcriptional activity. **B**. TFBSs scanned in accessible chromatin of *Obox6*. Expression levels of these TFs were high (FPKM ≥ 10). **C-F**. UCSC plots showing the epigenetic signal surrounding gene *Obox6* for 2-, 4-, and 8-cell embryos and ICM. **G**. Colormap showing the expression level of *Obox6* during preimplantation.

Here, we use the gene *Obox6* as an example to illustrate the relationship between different layers of epigenetic information and transcriptional activity (Fig. 6B-G). Soon after fertilization, *Obox6* was silenced; however, residual TF mRNAs from the oocyte and the zygote remained and corresponding TFBSs can be scanned in the promoter of *Obox6* (Fig. 6B), making it possible to open the chromatin. As expected, in the 2-cell stage, the promoter of *Obox6* was accessible, leading to active transcription of *Obox6*. At this time, the H3K4me3 signal was weak, and the mCG/CG level was high (Fig. 6C). In the 4-cell stage, chromatin accessibility of the gene promoter was sustained, while the H3K4me3 signal increased significantly; thus, *Obox6* was up-regulated (Fig. 6D). In the 8-cell embryo, expression of *Obox6* reached its highest level and a TAD boundary that *Obox6* located was constructed (Fig. 6E). Finally, in the ICM, as the promoter of *Obox6* was targeted with strong poised modifications including H3K27me3 and H3K9me3 but the chromatin accessibility and H3K4me3 signal decreased to a very low level, and gene expression was down-regulated (Fig. 6F).

## Discussion

### Extension to recent epigenomic landscape studies

Recent studies have provided the dynamic landscapes of different layers of epigenetic information, including chromatin accessibility, histone modifications, DNA methylation, and higher-order chromatin structure, in early human and mouse embryos [3, 9, 16, 21, 34, 35, 39–45]. Our work extends these studies by revealing how transcriptional programs combine with different epigenetic mechanisms in separate developmental periods. We liken the transcriptional program in early embryonic development to a drama, with the expectation that different “actors” help regulate gene expression in prescribed temporal and spatial contexts. Features of this drama include the opening of chromatin, resetting of histone modifications, condensing of the higher-order chromatin structure, and reprogramming of DNA methylation.

Chromatin accessibility represents TFBSs, many of which are located at enhancers [46]. We observed that chromatin accessibility is strongly associated with gene expression in earlier embryonic development (2- to 8-cell stages), whereas the association sharply decreases in the ICM and mESC stages (Fig. 1A). One possible explanation for this result is that early gene expression is merely dependent on the chromatin state at that time. Further regulation by H3K4me3 in the 4-cell stage and spatial regulation by the 3D chromatin structure depend on chromatin accessibility. However, with the addition of DNA methylation and poised histone modifications, such as histone H3 at lysine 27 trimethylation (H3K27me3), chromatin accessibility becomes progressively restricted and transcriptional activity becomes multi-regulated.

It is intriguing to speculate that accessible chromatin together with TF binding function as activators of transcription. However, we still do not completely understand what directs chromatin opening, nor do we know whether accessible chromatin establishes opening followed by selective closing or establishes selective opening from the onset. A previous study reported that pioneer TFs recognize closed chromatin regions and make them more accessible to the binding of other TFs [15]. Nyfa, for instance, promotes 2-cell DHS establishment and ZGA [16]. New technologies are needed to show genome-wide binding of pioneer TFs in early embryonic development.

### Chromatin accessibility contributes to the gain and loss of H3K4me3 in the 4-cell stage

It is of great interest to investigate the relationship between different epigenetic marks, previous studies have characterized the interplay between different epigenetic marks. For example, dimethylation of histone H3 at lysine 9 (H3K9me2) is positively correlated with DNA methylation [47], which, in turn, is negatively correlated with chromatin accessibility [48]. Monomethylation of histone H3 at lysine 4 (H3K4me1) is a predictor of chromatin loop interactions during stem cell differentiation [49]. However, to our knowledge, the relationship between H3K4me3 and chromatin accessibility during early embryonic development remains unknown. Although both chromatin accessibility and H3K4me3 are generally correlated with gene expression [16, 21, 50], whether they co-regulate transcriptional activity synchronously or asynchronously is not clear.

Our correlation analysis indicates that regulation of chromatin accessibility precedes H3K4me3 (Fig. 1A and Fig. 3A). Chromatin accessibility increases sharply in the 2-cell stage and gradually decreases thereafter (Fig. 1B), whereas the H3K4me3 level increases significantly in the 4-cell stage (Fig. 3C). H3K4me3 peaks are depleted in zygotes and can be observed after major ZGA [9], indicating the completion of H3K4me3 reprogramming in late 2-cell embryos, which is later than the previously reported rapid reprogramming of chromatin accessibility at the PN3 stage [16]. As shown in Fig. 3D and Fig. 3F, the gain and loss of H3K4me3 peaks are accompanied by the opening and closing of chromatin. We speculate that TFs recruit “writer” complexes, such as SET1, MLL1, and MLL2 [51, 52], for H3K4me3 by binding to accessible chromatin, thereby activating H3K4me3 to regulate gene expression. Experiments are required to validate whether accessible chromatin is necessary for H3K4me3 to regulate gene expression. Collectively, the findings from our study not only describe the interplay between different layers of epigenetic information but also provide deeper insights into the relationship between these markers and gene expression in a spatiotemporal manner, helping to elucidate the fundamental mechanisms of transcriptional programs during early embryonic development.

## Meterials and Methods

### Data

ATAC-seq data for mouse early 2-cell, 2-cell, 4-cell, and 8-cell stages and ICM were obtained from GSE66390. DHS assay data for mouse oocyte, sperm, 1-cell, and 2-cell stages were obtained from GSE76642. ChIP-seq data for TFs in human H1 stem cells were obtained the ENCODE project. Data for histone modification H3K4me3 for mouse early 2-cell, 2-cell, 4-cell, and 8-cell stages and ICM were obtained from GSE71434. Hi-C data for mouse early 2-cell, 2-cell, and 8-cell stages, ICM, and mESC were obtained from GSE82185. DNA methylation data for mouse 2-cell and 4-cell stages and ICM were obtained from GSE56697 and for mouse E3.5TE, E4.0ICM, E5.5Epi, E5.5VE, E6.5Epi, and E6.5VE and germ layers were obtained from GSE76505. Mouse gene annotations were obtained from Mouse Genome Informatics (MGI) [53]. Our study was based on the mm9 genome.

### Density of epigenetic information in gene promoter

For chromatin accessibility and H3K4me3, we first counted the total ATAC-seq or ChIP-seq reads at the gene promoters (2 kb upstream of the TSSs) and then called the density by calculating the FPKM of each marker. For DNA methylation, because MethylC-Seq was aligned to mm10, we first counted the total CG reads and methylated CG reads at the gene promoter (2 kb upstream of the TSS) based on the mm10 genome, and then quantified the methylation level by calculating the mCG/CG level.

### Correlation between gene expression and epigenetic information

For each developmental stage, we sorted genes according to the descending level of gene expression, divided genes into 100 copies, and calculated a mean value for each copy. Correlation coefficients (R^2^ values) and *p*-values were calculated by Pearson’s linear correlation.

### Dynamics of accessible chromatin during preimplantation

For Supplementary Fig. 1A, accessible chromatin that was “maintained” in current stage was defined as accessible chromatin belonging to both current stage and previous stage (using *bedops –e −25%*). Accessible chromatin that was “gained” in current stage was defined as accessible chromatin belonging to current stage but not found within previous stage (using *bedops –n −25%*) and accessible chromatin that was “lost” in previous stage was defined as accessible chromatin belonging to previous stage but not found within current stage (using *bedops –n −25%*).

### Identification of TFBSs

Position-specific weight matrices (PWMs) of 395 TFs, which corresponded to 427 motifs, were collected from the HOCOMOCO (v10) database [54]. Genomic sequences from an accessible chromatin region in the mm9 genome were used as input for FIMO [55]. A custom library containing all 427 motifs was used to scan for motifs at a *p*-value threshold of 10^−5^. For each TF, multiple TF motifs, if present, were combined to generate the corresponding TFBSs.

### Identification of significant interactions and TAD boundaries

Hi-C data from previous work [56] were collected and subjected to the HOMER [57] algorithm to identify significant interactions with a default parameter (*p*-value = 0.001) and 5-kb resolution. A TAD was detected by using a 40-kb resolution normalized contact matrix described in previous work [58]. TAD boundaries were identified by using the insulation score method at 40-kb resolution [59], with two minor modifications. First, a 200-kb genomic region, rather than a 500-kb region, was used to detect TAD (which is now considered to be mostly under 200 kb). Second, a 120-kb window, rather than a 100-kb window, was used to fit the resolution in delta vector calculations.

**Author contributions**

**Acknowledgments**

## SUPPLEMENTARY MATERIALS

Supplementary Fig. 1. General characterization of accessible chromatin during early embryonic development.

Supplementary Fig. 2. The 4-cell stage: H3K4me3 facilitates gene up-regulation.

Supplementary Fig. 3. 3D chromatin structures enhance transcriptional activity in the 8-cell stage.

Supplementary Fig. 4. Multifaceted of epigenetic marks at ICMs.

Supplementary Fig. 5. Summary of active genes and silence genes at ICMs.

Supplementary Fig. 6. Different layers of epigenetic marks shape transcriptional programs.

